# Visual Field Analysis: a reliable method to score left- and right eye-use using automated tracking

**DOI:** 10.1101/2021.05.08.443242

**Authors:** Mathilde Josserand, Orsola Rosa-Salva, Elisabetta Versace, Bastien S. Lemaire

**Author notes:** author for correspondence: Mathilde Josserand & Bastien S. Lemaire.

## Abstract

Brain and behavioural asymmetries have been documented in various taxa. Many of these asymmetries involve preferential left- and right-eye use. However, measuring eye use through manual frame-by-frame analyses from video recordings is laborious and may lead to biases. Recent progress in technology allowed the development of accurate tracking techniques for measuring animal behaviour. Amongst these techniques, DeepLabCut, a python-based tracking toolbox using transfer learning with deep neural networks, offers the possibility to track different body parts with unprecedented accuracy. Exploiting the potentialities of DeepLabCut, we developed ‘Visual Field Analysis’, an additional open-source application for extracting eye-use data. To our knowledge, this is the first application that can automatically quantify left-right preferences in eye use. Here we test the performance of our application in measuring preferential eye-use in young domestic chicks. The comparison with manual scoring methods revealed a perfect correlation in the measures of eye-use obtained by ‘Visual Field Analysis’. With our application, eye-use can be analysed reliably, objectively and at a fine scale in different experimental paradigms.

## Introduction

Quantifying accurately animal behaviour is crucial to understand its underlying mechanisms. Historically, behavioural measurements were collected manually. However, with technological progress, automated data collection and analyses have expanded (Anderson and Perona 2014), making behavioural analyses more precise, reliable and effortless for the experimenter (Wood and Wood 2019; Versace et al. 2020; Lemaire et al. 2021),

Automated data collection may allow finer and more objective behavioural analyses than the ones provided by manual coding. However, many of the currently available tracking techniques and software can be complex to use and even inaccurate in some experimental conditions (such as in poor and changing illumination, low contrasts, etc.). The open-source toolbox DeepLabCut copes with these limitations (Mathis et al. 2018; Nath et al. 2019). DeepLabCut exploits deep learning techniques to track animals’ movements with unprecedented accuracy, without the need to apply any marker on the body of the animal (Labuguen et al. 2019; Wu et al. 2019; Worley et al. 2019), opening a new range of possibilities for measuring animal behaviour such as preferential eye-use.

It is now clear that structural and functional asymmetries, once believed to be unique of humans, are widespread among vertebrates (Rogers et al. 2013; Versace and Vallortigara 2015; Vallortigara and Versace 2017) and invertebrates (Frasnelli et al. 2012). The study of sensory and perceptual asymmetries is a powerful tool to understand functional lateralization, especially in animals with laterally placed eyes. For instance, in non-mammalian models, researchers can take advantage of anatomical features causing most of the information coming from each eye-system to be processed by the contralateral brain hemisphere (Vallortigara and Versace 2017). To date, temporary occlusion of one eye has been the main method used for behavioural investigation of these eye asymmetries (Güntürkün 1985; Vallortigara 1992; Vallortigara et al. 1999; Chiandetti 2017), However, studying the spontaneous eye-use without monocular occlusions is important to shed light on the lateralization of naturalistic behaviours. Indeed, a considerable amount of evidence has shown that animals actively use one or the other visual hemifield depending on the task (Güntürkün and Kesch 1987; Vallortigara et al. 1996; Sovrano et al. 1999; Tommasi et al. 2000; De Santi et al. 2001; Santi et al. 2002; Prior et al. 2004; Rogers 2014; Schnell et al. 2018) and/or motivational/emotional state (Andrew 1983; Bisazza et al. 1998).

A widespread method to test preferences in eye use is the frame-by-frame analyses of video recordings (Fagot et al. 1997; Rogers 2019). A drawback of this procedure is that it is tedious and may originate potential errors and biases (Anderson and Perona 2014). Moreover, in animals with two foveae (or a ramped fovea) like birds, it is necessary to distinguish the use of frontal and lateral visual fields, making manual coding of these data even more complicated (Vallortigara et al. 2001; Lemaire et al. 2019). To address these issues, we developed an application for the automatic recording of eye-use preferences for comparative neuroethological research. This application, named ‘Visual Field Analysis’, is based on DeepLabCut tracking (Nath et al. 2019) and enables eye-use scoring as well as other behavioural measurements (see Josserand and Lemaire 2020 for more details).

Our application can be used to measure eye-use for any setting in which one animal is video-recorded from the top by a fixed camera. It aims to measure eye-use preferences while observing different stimuli (an approach that has been used, for instance, in Dharmaretnam and Andrew 1994; De Santi et al. 2001; Vallortigara et al. 2001; Rogers et al. 2004; Dadda and Bisazza 2016; Schnell et al. 2016). Moreover, it is possible to measure other variables such as the level of locomotor activity (a measurement used in open field or runway tests, e.g. Gallup and Suarez 1980; Gould et al. 2009; Ogura and Matsushima 2011) and how much time the animal spends in different areas of the test arena (a measure widely used in recognition, generalization and spontaneous preference tests, which measure animals’ preferences between two stimuli, Wood 2013; Rosa-Salva et al. 2016; Versace et al. 2016). Using Visual Field Analysis, eye-use can be assessed when the animal is moving or still and while it is looking at one or two stimuli. These can also be either stationary or in motion, but must be placed at opposite sides of the apparatus. Eye-use is calculated as the number of frames spent looking at a stimulus with either one eye or the other. Furthermore, each hemifield can be divided into two separate visual fields (frontal and lateral, of adjustable size). This is particularly relevant for species that use their frontal and lateral visual fields differently (e.g., domestic chicks and king penguins, Vallortigara et al. 2001; Lemaire et al. 2019). Moreover, Visual Field Analysis measures the movements of the animal’s head, by reporting the number of pixels crossed by the tracking-point associated with the top of the animal’s head over several frames. This measure can reveal the overall locomotor activity of an animal moving freely in its environment or the extent of the head movements of a restrained animal. In this latter case, our activity measurement can inform about head saccades, a behaviour present in birds, which is correlated with arousal (Kjrsgaard et al. 2008; Golüke et al. 2019). Finally, the time spent by an animal in up to five different areas of a rectangular arena can be conviently monitored by Visual Field Analysis. For further details on the functioning and current limitations of this application, please see the full protocol published by Josserand and Lemaire (2020).

The aim of the current study was to provide an experimental validation of the main function of Visual Field Analysis: scoring preferential eye-use in animals with laterally placed eyes. To do so, we assesed the accuracy of Visual Field Analysis in scoring preferential eye use of domestic chicks (*Gallus gallus*), while looking at an unfamiliar stimulus, in comparison with traditional manual scoring.

## Methods

### Subjects

The experimental procedures were approved by the Ethical Committee of the University of Trento and licenced by the Italian Health Ministry (permit number 53/2020). We used 10 chicks, of undetermined sex (strain Ross 308). The eggs were obtained from a commercial hatchery (Azienda Agricola Crescienti) and incubated at the university of Trento under controlled conditions (37.7°C and 40% of humidity). Three days before hatching, we moved the eggs into an hatching chamber (37.7°C and 60% of humidity). Soon after hatching, the chicks were housed together within a rectangular cage (150 x 80 x 40 cm) in standard environmental conditions (30°C and homogenous illumination, adjusted to follow a natural day/night cycle) and in groups of maximum 40 individuals. Food and water were available ad libitum. The animals were maintained in these conditions for 3 days, until the test was performed. After the test, all animals were donated to local farmers.

### Test

The test took place the third day post-hatching. Each chick was moved into an adjacent room and placed in a smaller experimental cage (45 x 20 x 30 cm) to begin the pre-test habituation phase, which usually lasted about 30 minutes. The cage had a round opening (4cm), and during the habituation phase the animal could pass its head trough it at will, to inspect an additional empty compartment (20 x 20 x 30 cm, see Figure 1). Young chicks tend to spontaneously perform this behaviour, when given the opportunity. Once a subject was confidently passing its head through the opening, the proper test phase began and a red cylinder (5 cm high, 2 cm diameter) was added in the additional compartment (20 cm away). The subject’s head was then gently placed through the round opening by the experimenter and and the animal was manually kept in this position for 30 seconds (Figure 1). The behaviour of each animal was recorded with an overhead camera (GoPro Hero 5, 1290×720, ∼ 25 - 30 fps) for 30 seconds. Each animal was tested only once.

**Figure 1:**
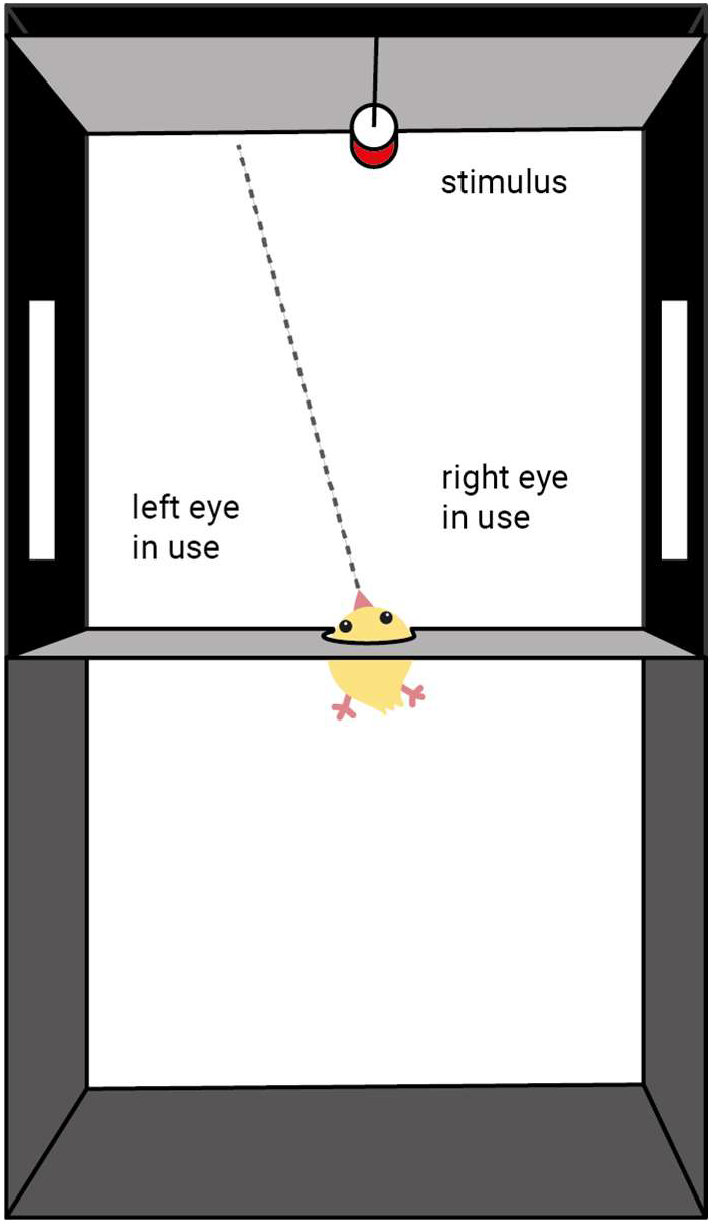
Schematic representation of the testing condition for Experiment 1 (top view).

### Data acquisition using Visual Field Analysis

#### Data acquisition

To perform data acquisition, Visual Field Analysis requires three main inputs for each analysed subject (Figure 2). The first input is the video recording of the animal’s behaviour. The second input is a file containing information about the tracking (x, y coordinates) of specific body parts (output file provided by DeepLabCut). ‘Visual Field Analysys’ focuses on three points located on the head: the closest points to the left eye, the right eye and the top of the head. For the current experiment, the positions of these 3 points were manually labelled on 100 frames so that DeepLabCut could accurately generalise each point of interest on all video recordings. The third input corresponds to a spreadsheet where the experimenter manually enters specific information about the observed animal. Further information is provided in our protocol (Josserand and Lemaire 2020).

**Figure 2:**
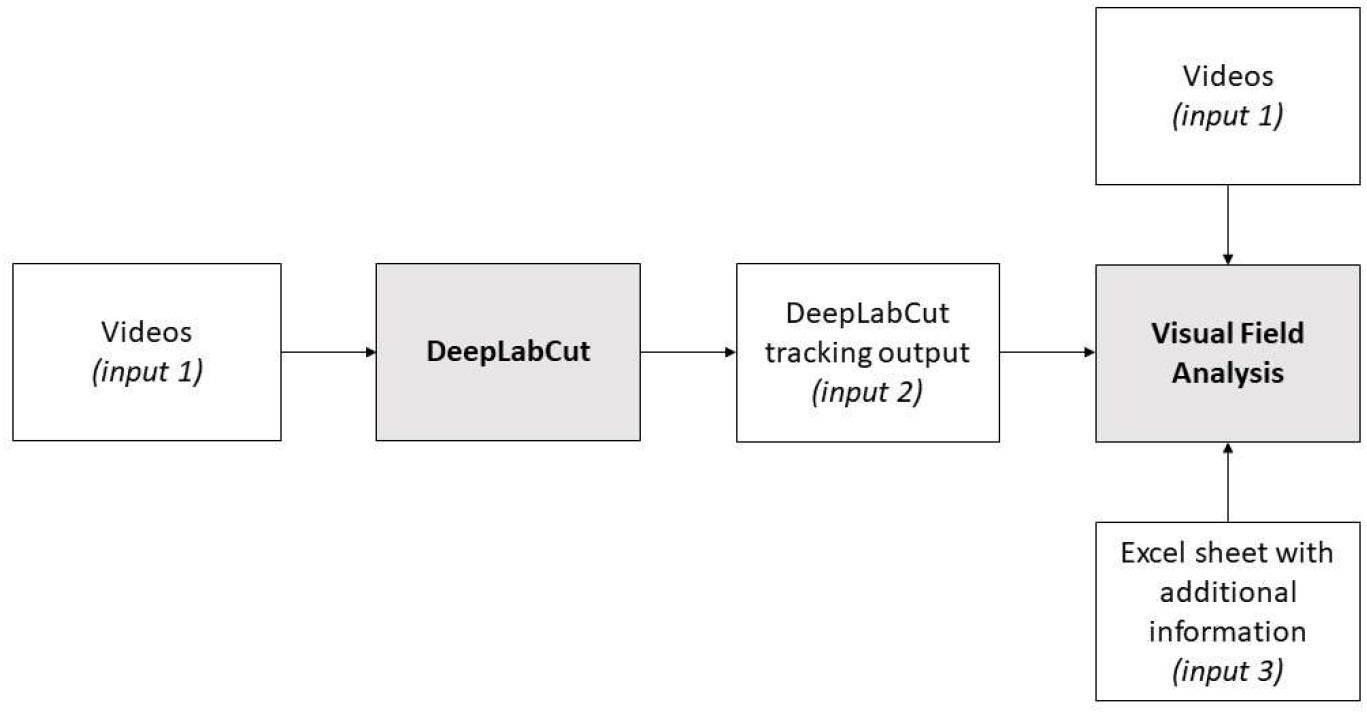
Diagram showing the inputs required to run Visual Field Analysis.

To proceed with data acquisition, the extent of the frontal and lateral visual fields for the species under investigation must also be defined. In the current study, we subdivided the visual field into: two frontal visual fields (each 15° wide from the midline, Figure 3), two lateral visual fields (each 135° wide starting from the frontal visual field line, Figure 3) and the blind visual field (30° wide starting from the lateral visual field line, Figure 3).

**Figure 3:**
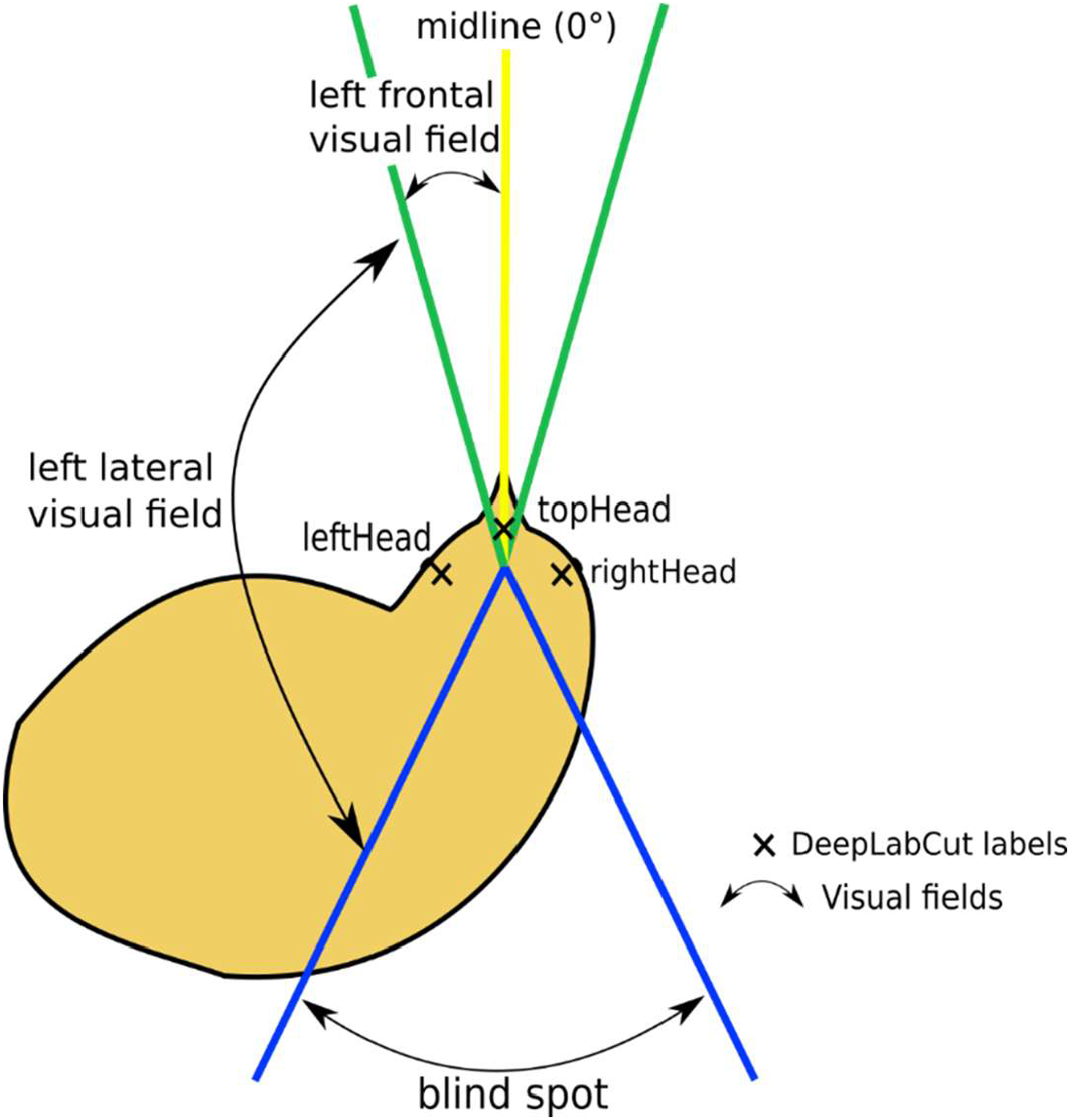
Schematic representation of the visual fields defined for the experiments. The yellow midline separates the left visual field from the right visual field. The green lines show the borders of the frontal vision (from the midline, 15° on each side) and the blue lines show the blindspot of the chick. Each angle can be manually chosen in Visual Field Analysis program.

Using the visual fields previously defined and the location of the stimuli, the program assesses in which portion (frontal or lateral) of the hemifield (left or right) the stimuli fall in each frame. For each frame, if the stimulus(i) is located within a visual field, a value of 1 is attributed to that visual field (see the light-green dash line on Figure 4.A). If the stimulus is straddling on two visual fields, the proportion of the object located within each visual field is attributed to each one of them (see the light-green dash line on Figure 4.B).

**Figure 4:**
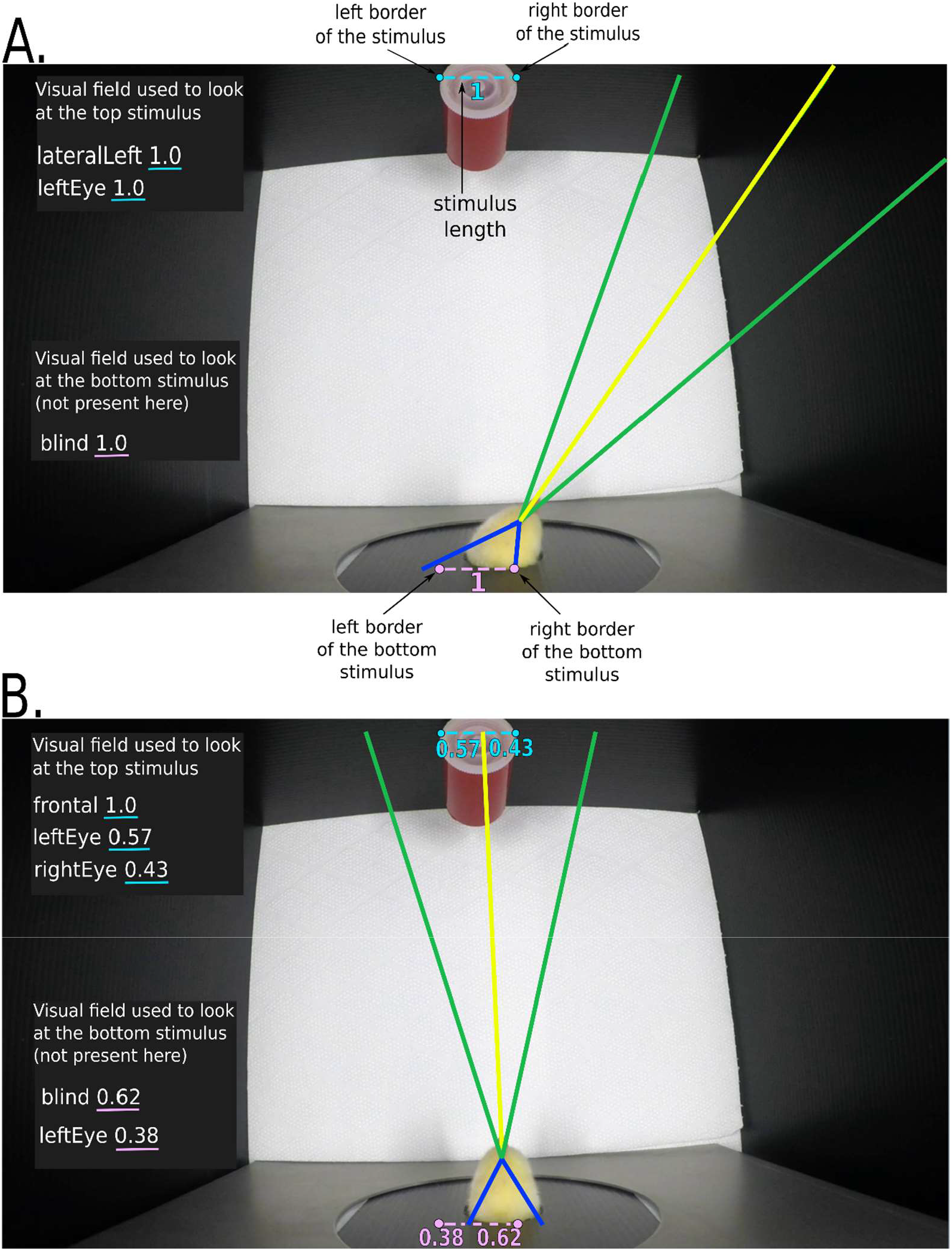
Visualisation of the projection lines defining each region of the visual field. The yellow line indicates the midline and delimitates each hemifield, providing the nasal margin of the frontal visual fields. The green lines delimitate the frontal visual fields from the lateral visual fields. The blue lines delimitate the lateral visual fields from the blind spot. In these pictures, referring to the set up of the current experiment, the visual field used to look at the stimulus is shown, which can be compare to the information reported on the left side of the pictures (within the dark rectangles on top and bottom left corners of the images). Since our application allows to record visual field use for up to two simultaneously presented objects, in these images we can see two grey rectangles (reporting information on the eye-use for each of the two objects). In the current example, where only one stimuls was present, the relevant information is presented in the grey rectangle in the upper part of the image (referring to the top stimulus), while the other can be ignored. A value is assigned to every visual field, indicating whether the stimulus was located inside it. A value of 1 for a given visual field indicates that the stimulus is entirely located within that visual field, such as in figure 4A. However, the stimulus can be straddling into two visual fields, such as in figure 4B. Consequently, the program attributes different values depending on the portion of the stimuli extent (lillac dash lines) located in a visual field (the stimulus extent is here defined by its borders).

The output through which ‘Visual Field Analysis’ provided eye-use data varies depending on the location and number of stimuli. In this experiment, Visual Field Analysis computed eye-use for one stimulus.

#### Error threshold setting

As a strategy to identify and exclude frames innacurately tracked by DeepLabCut, we implemented an error threshold approach. For each video the experimenter can set an error threshold that specified the acceptable range of distances between the three points tracked on the head of the animal. The error threshold is based on the average distance between each body parts tracked by DeepLabCut and excludes frames which are too far from the average distance. As an example, the average distance between the ‘leftHead’ and ‘rightHead’ points was 58 pixels for subject 5 of the current experiment. For this animal, we chose a threshold of 3. This threshold corresponds to the number of standard deviations above which a frame is considered as an outlier; thus, the absolute value of the threshold in terms of pixels is variable. With a threshold set at 3, 1.86% of the frames were preliminarly labelled as outliers. These frames were then manually inspected to address the accuracy of this process and could be removed from our analysis if visual observation confirmed them to be outliers (Figure 5). The treshold used for each subject of the current experiment, as well as the percentage of frames manually checked and excluded from the analyses at these different steps, are reported in Table 1.

**Figure 5:**
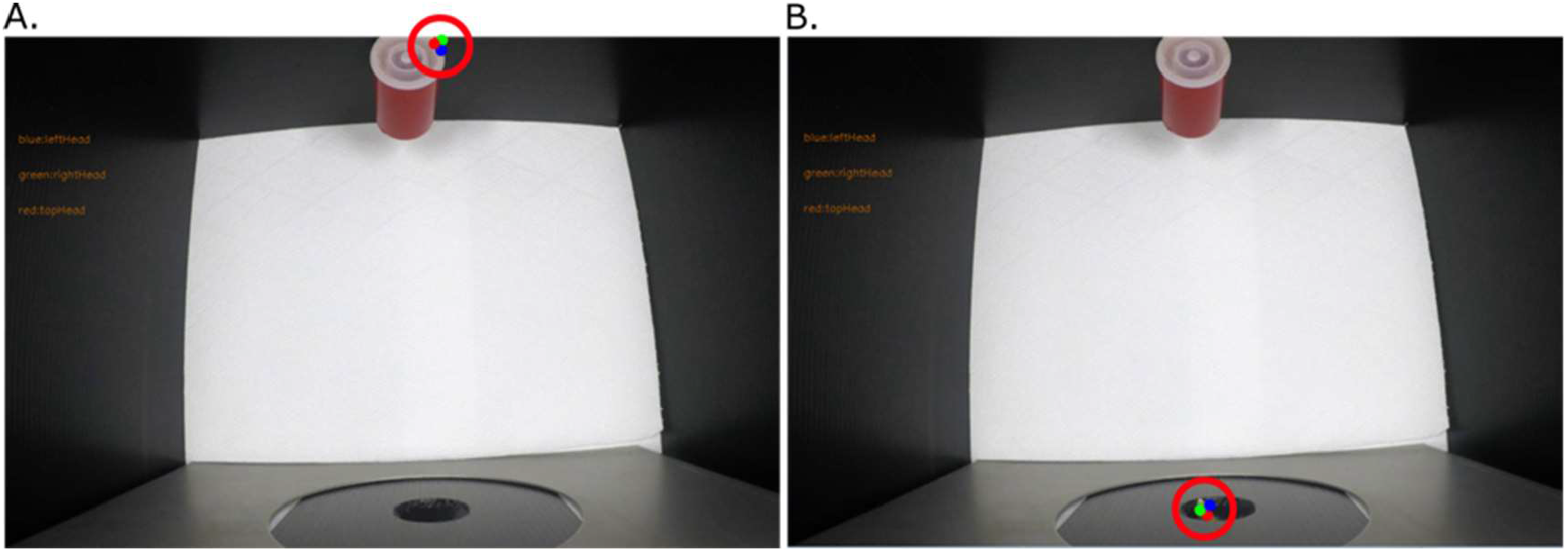
Two of the frames considered as outliers in our example (from subject 5 of the current experiment), with a threshold of 3. The red circles on the images highlights the position of the labels. On image 5A, the chick has not placed its head inside the round opening yet, but DeepLabCut incorrectly placed the ‘leftHead’ (blue dot), ‘topHead’ (green dot) and ‘rightHead’ (red dot) on an empty portion of the screen, close to the stimulus. On image 5B, the chick started to insert its head in the round opening, but most of it is still invisible. DeepLabCut incorrectly located the ‘leftHead’, ‘topHead’ and ‘rightHead’ labels on the animal’s beak.

**Table 1:**
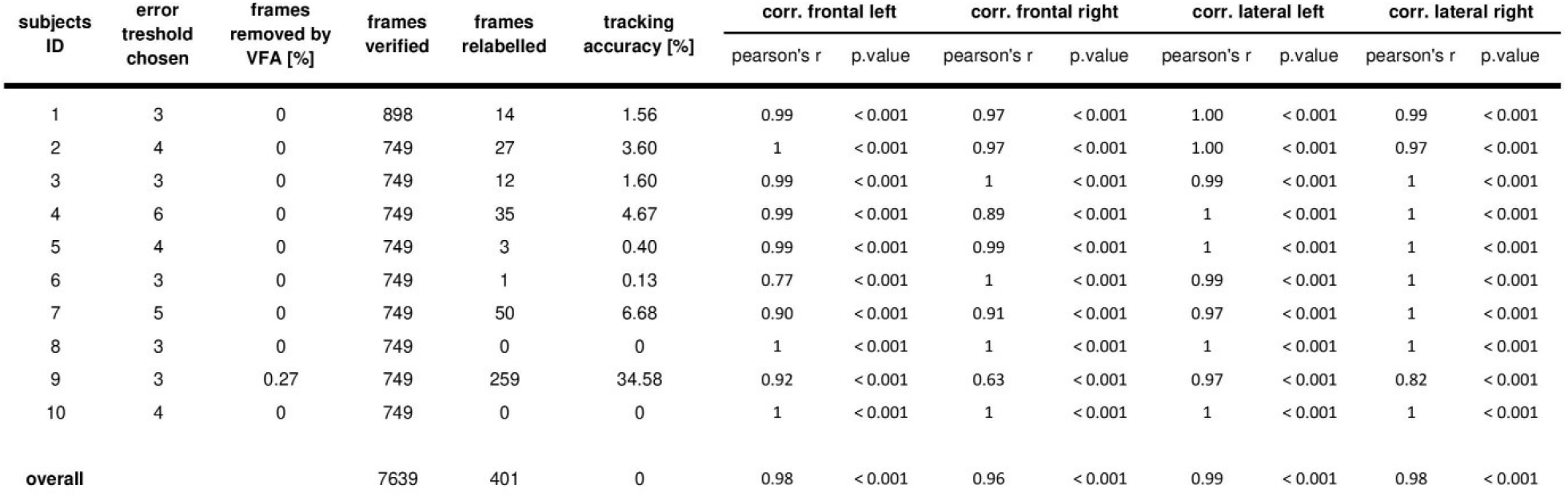
Table showing the tracking accuracy at the individual and at the group level. All the frames analysed by Visual Field Analysis were then manually coded, and the “ frames relabelled” column reports the number of frames for which a discrepancy emerged between manual and automated coding. The “ frame verified” column reports the tracking accuracy, which correspond to the number of frames correctly tracked (“ frames verified” minus “ frames relabelled”) over the total number of frames (“ frames verified”). The two types of coding (manual and automatic) have been compared using Spearman’s correlation tests, both at the individual level and for the whole sample of N=10 chicks. This has been done separately for each visual field (frontal left and right, lateral left and right). Results of these correlation are reported in the last 8 columns.

| subjects ID | error threshold chosen | frames removed by VFA [%] | frames verified | frames relabelled | tracking accuracy [%] | corr. frontal left |  | corr. frontal right |  | corr. lateral left |  | corr. lateral right |  |
| --- | --- | --- | --- | --- | --- | --- | --- | --- | --- | --- | --- | --- | --- |
|  |  |  |  |  |  | pearson's r | p.value | pearson's r | p.value | pearson's r | p.value | pearson's r | p.value |
| 1 | 3 | 0 | 898 | 14 | 1.56 | 0.99 | < 0.001 | 0.97 | < 0.001 | 1.00 | < 0.001 | 0.99 | < 0.001 |
| 2 | 4 | 0 | 749 | 27 | 3.60 | 1 | < 0.001 | 0.97 | < 0.001 | 1.00 | < 0.001 | 0.97 | < 0.001 |
| 3 | 3 | 0 | 749 | 12 | 1.60 | 0.99 | < 0.001 | 1 | < 0.001 | 0.99 | < 0.001 | 1 | < 0.001 |
| 4 | 6 | 0 | 749 | 35 | 4.67 | 0.99 | < 0.001 | 0.89 | < 0.001 | 1 | < 0.001 | 1 | < 0.001 |
| 5 | 4 | 0 | 749 | 3 | 0.40 | 0.99 | < 0.001 | 0.99 | < 0.001 | 1 | < 0.001 | 1 | < 0.001 |
| 6 | 3 | 0 | 749 | 1 | 0.13 | 0.77 | < 0.001 | 1 | < 0.001 | 0.99 | < 0.001 | 1 | < 0.001 |
| 7 | 5 | 0 | 749 | 50 | 6.68 | 0.90 | < 0.001 | 0.91 | < 0.001 | 0.97 | < 0.001 | 1 | < 0.001 |
| 8 | 3 | 0 | 749 | 0 | 0 | 1 | < 0.001 | 1 | < 0.001 | 1 | < 0.001 | 1 | < 0.001 |
| 9 | 3 | 0.27 | 749 | 259 | 34.58 | 0.92 | < 0.001 | 0.63 | < 0.001 | 0.97 | < 0.001 | 0.82 | < 0.001 |
| 10 | 4 | 0 | 749 | 0 | 0 | 1 | < 0.001 | 1 | < 0.001 | 1 | < 0.001 | 1 | < 0.001 |
| overall |  |  | 7639 | 401 | 0 | 0.98 | < 0.001 | 0.96 | < 0.001 | 0.99 | < 0.001 | 0.98 | < 0.001 |

To help users choosing an appropriate threshold and check the program’s accuracy, Visual Field Analysis has a built-in function to visualise frames. It is possible to either visualize the frames removed from the analysis given the chosen error threshold, in addition to a set of randomly selected frames wich are provided by the program to assess its accuracy.

#### Manual coding of the data

To assess the reliability of the eye-use data provided by Visual Field Analysis, we manually checked all the frames analysed by Visual Field Analyses (7639 frames in total). The frames were checked and saved using a built-in function provided by Visual Field Analysis. Each frame was inspected by two independent coders, each of whom coded all the frames independently from the other. Then, the two experimenters compared their output, re-inspected all the frames for which a different coding was assigned and agreed on a final labelling of these frames.

The manual scores were attributed using the same score attribution method and the same visual field subdivisions than Visual Field Analysis. On each frame, the coders surimposed a transparent sheet, on which five lines originating from a central point represented the subdivisions of the chick visual field (see Fig. 3). For each frame, if the visual-field delimitation lines of the translucent sheet used for manual scoring overlapped perfectly with the lines used by Visual Field Analysis for the attribution of the scoring (as visible in Fig. 4), the original automated scoring was considered accurate. In this case, the same original scoring provided by Visual Field Analysis was reported also for manual scoring. If there was not a perfect overlap, the original scoring was considered inaccurate and the frame was “ relabelled”, providing new vlaues for manual scoring. In this case, the same criteria described above were followed: if the stimulus entirely fallen out within a visual field, a value of 1 was attributed to it. If the stimulus straddled on two visual fields, the proportion of the stimulus located within each visual field was attributed to each one of them (e.g., 0.75 and 0.25).

### Statistical analyses

The scores obtained for each visual field (frontal left, frontal right, lateral left and lateral right) were compared between scoring methods (Visual Field Analysis vs. manual coding) using correlation tests (Pearson’s test) overall and per individual. As some videos were better tracked by DeepLabCut than others, we report the reliability of our program in relation to the tracking accuracy (measured as the percentage of frames that received identical scoring in the manual and automated scoring). The statistical tests were performed using RStudio (RStudio Team 2015).

## Results

### Eye-use data reliability

The Pearson’s correlations tests revealed an almost perfect, and highly significant, correlation between scoring methods, both when the data of the whole sample were taken in in consideration as well as when the analysis was run at the single subject level. This was true for each visual field (see Table 1 for statistics).

The number of frames for which the manual scoring of the human coder was discrepant from that assigned by Visual Field Analysis (relabelled frames) is detailed for each subject in Table 1. No discrepancy between the manual coding and the automated coding was found for subject 8 and 10. Thus, for these subjects, no frames were relabelled and a perfect correlation was, of course, found between manual and automated scoring. One should note that, the reliability of our application is directly dependent on the DeepLabCut tracking accuracy, which differs across conditions (different videos settings) and individuals (different behaviours). When the DeepLabCut tracking was 100% accurate, the output produced by Visual Field Analysis perfectly matched the manual scoring done by the human coders.

For the 8 remaining subjects, the tracking accuracy (i. e. the percentage of frames that received identical scoring in the manual and automated scoring) fluctuated from 65.4 to 99.9%. Nonetheless, the reliability of the program to score eye-use remained relatively high in all conditions (see Table 1 for statistics). Even when the tracking accurary was at its lowest (65.4 % for subject 9), the correlation between the coding provided by Visual Field Analysis and the manual coding remained strong for most visual fields (pearson’s r ranging from 0.82 to 0.97), although it decreased in the frontal right visual field (Pearson’s r = 0.63).

## Discussion

Automated and reliable assessmets of visual field use can support the investigation of behavioural lateralisation. Our results show that Visual Field Analysis can reliably be used to automatically assess eye-use behaviour in animals with laterally-placed eyes, substituting manual coding methods. The comparison between the manual and of the automaned scoring revealed a nearly perfect correlation between manual score and VIdual Field Analysis score.

With optimum tracking conditions, the results provided by the application can be reliable at 100% (i.e., identical results can be obtained as with manual scoring). Visual Field Analysis excludes from the analyses frames with a low degree of likelihood (i.e., frames that DeepLabCut considered as being unlikely to be well-tracked), thus keeping only frames with a confidence of being well-tracked above 95%. It also excludes frames where the distance between the DeepLabCut’s labels is higher than a given threshold. Note that the two measures are often correlated: when DeepLabCut tracking is imperfect, both the number of frames considered as unlikely to be well-tracked and the number of frames considered as outliers with the threshold method are high. Therefore, most of the frames that could be wrongly tracked are excluded from the analysis. Moreover, unsing a built-in function of our application, the user can manually visualize a certain number of random frames of a video to check the program performance. We suggest to visualise at least 100 frames per individual with more than 90% of tracking accuracy, in order to achieve a performance similar to what described in the current study If the tracking accuracy is lower than 90%, we suggest to train again DeepLabCut using a new set of frames or a different labelling method.

Visual Field Analysis can not only assess simple preferential eye-use (whether the left or right eye is preferred). It can also investigate the use of sub-regions within each hemifield (frontal visual field vs lateral visual field), allowing the analysis of eye-use behaviour at a fine level, which would be very time consuming if done manually. Alongside measuring eye-use, the application allows the recording of other relevant behavioural measurements, such as the activity level of an animal’s head while keeping track of its positions in different areas of a testing environment (for this last function, the performance of Visual Field Analysis has been already validated in a previous study, showing once again very high correlation with the measurements obtrained by traditional manual coding methods (Santolin et al. 2020)). This provides additional information which can be analysed in relation to eye-use behaviour or independently from it, allowing for richer behavioural assessements and for more flexible use in different experimental designs.

Given the numerous practical advantages offered by automated behavior tracking methods, compared to manual ones, we believe that automated methods should be chosen to ensure reproducible data analysis. Visual Field Analysis offers an important resource for research on behavioural lateralization, allowing to collect and analyse a richer set of data, in a less time consuming and unbiased way.

## Acknowledgements

We thank Anastasia Morandi-Raikova for helping with the manual coding of the frames. We also thank Prof. Giorgio Vallortigara for funding MJ during her stay at CIMeC, and for his support for this project. Finally, we are also thankful to Lise Lecroq for sharing with us the Illustration of Figure 1.

## Author Contributions

BL implemented DeepLabCut. MJ designed and coded VFA. BL did part of the manual coding. EV contributed to the experimental design, and suggested ways to validate VFA tool. BL wrote first draft with critical edits from ORS and EV. All authors contributed to the manuscript.

## Open Practices Statement

‘Visual Field Analysis’ is open-source and entirely available in our <u>GitHub</u>. Videos and data are available on fig**Share** (https://figshare.com/s/ecfc65e0bf4d562a0fc5). A step by step guide tu run our program is available on protocols.io (Josserand and Lemaire 2020).

